# The Effects of Sex and Strain on *Pneumocystis murina* Fungal Burdens in Mice

**DOI:** 10.1101/781245

**Authors:** Nikeya L. Tisdale, Alan Ashbaugh, Keeley Hendrix, Margaret S. Collins, Alexey P. Porollo, Melanie T. Cushion

**Affiliations:** University of Cincinnati College of Medicine, Department of Internal Medicine, Division of Infectious Diseases, Cincinnati, OH 45267; Cincinnati VAMC, Medical Research Service, Cincinnati, OH 45220; Center for Autoimmune Genomics and Etiology and Division of Biomedical Informatics, Cincinnati Children’s Hospital Medical Center, Cincinnati, OH 45229, USA

**Keywords:** Pneumocystis, pneumonia, AIDS-related, sex differences, experimental design

## Abstract

Many preclinical studies of infectious diseases have neglected experimental designs that evaluate potential differences related to sex with a concomitant over-reliance on male model systems. Hence, the NIH implemented a monitoring system for sex inclusion in preclinical studies.

**Methods:** Per this mandate, we examined the lung burdens of *Pneumocystis murina* infection in 3 mouse strains in both male and female animals at early, mid and late time points.

**Results:** Females in each strain had higher infection burdens compared to males at the later time points.

**Conclusion:** Females should be included in experimental models studying Pneumocystis.

## 1. Introduction

The National Institute of Health Revitalization Act of 1993 mandated the inclusion of women participants in federally funded clinical research. With this inclusion of female participants, clinical research findings began to show differences in the drug response outcomes in women compared to men participants [1]. However, the use of females in experimental model systems was not widely adopted in preclinical research studies. A 2011 report noted the underrepresentation of female animals in 8 out of 10 biological disciplines [2]. Subsequently, on October 2014, the NIH implemented a monitoring system for sex inclusion in preclinical research, mandating federally funded researchers to report plans of the use of both male and female cell lines and animal models in preclinical studies or to explain why this was not necessary [3].

It is generally held that females develop more robust innate and adaptive immune responses than males and are less susceptible to infections caused by bacteria viruses, parasites and fungi [4]. However, there are exceptions to this rule and in some cases, contradictory results that were clearly dependent on the experimental model. For example, in a model of murine cutaneous leishmaniasis, males were more resistant to a physiological low-dose infection but more susceptible to higher doses and other methods of inoculation [5]. In another example, male and female mice differed in their responses to infection with *Plasmodium berghei*; whereas both sexes experienced peak parasitemia at the same time and time to death was the same, weight loss and brain responses were increased in males vs females [6].

In the present report, we focused on sex differences associated with Pneumocystis pneumonia (PCP), an infection caused by pathogenic yeast-like fungi in the genus Pneumocystis. In humans, PCP infects males and females with compromised immune systems and can cause life-threatening pneumonia. Strikingly, there was but a single publication in the PubMed database that compared PCP outcomes in men and women with HIV in 1987 showing women were more likely than men to die in the hospital from PCP [7]. A more recent meta-analysis showed an increase of PCP in non-immunodeficiency virus (HIV) infected patients, where being a female non-HIV patient was a risk factor, resulting in an increased mortality rate in female vs male non-HIV patients [8].

Investigators have generally used male rodents for studies of Pneumocystis infection, especially for preclinical drug development [9]. The identification of new prophylactic and therapeutic anti-Pneumocystis agents is a critical focus of investigation as there are few drugs with which to treat PCP. It is not known whether there are differences in responses by the sexes to this infection, but it could be an important consideration in future experimental designs. In this report, we compared the fungal burdens of *P. murina* in dexamethasone-immunosuppressed male and female mice to address a fundamental aspect of the mouse model of infection.

## 2. Material and Methods

### Design

Corticosteroid-immunosuppressed male and female Balb/c, C3H/HeN, and C57BL/6 mice (Charles River Laboratories) were infected with *P. murina* through exposure to *P. murina-*infected and immunosuppressed mice (seed mice) of the same strain and gender, following previous protocols [10, 11]. Seed mice have existing fulminate infections and transmit the infection to the naïve immunosuppressed mice through an airborne method of transmission [10]. The recipient (and seed mice) mice were immunosuppressed by adding dexamethasone at 4µg/liter to their drinking water and housed with the seed mice for 2 weeks. The mice were fed autoclaved standard lab chow from Charles River Laboratories (Wilmington, MA) and administered acidified water (1MHCL) to discourage secondary microbial infections, *ad libitum*. At 4-, 6-, and 8-week post-exposure, 8 male and female mice of each strain were sacrificed. Asci and total nuclei were enumerated as describe[12]. Microscopic enumeration of asci assesses the numbers of this life cycle stage which is alleged to be the product of the sexual cycle of these fungi as well as its transmissive stage. Enumeration of nuclei represents total fungal burden as the nuclei of all life cycle stages are counted.

### Evaluation of organism burden

At each of the time points, mice were euthanized by carbon dioxide inhalation per IACUC approved methods and the lungs were removed. After dissection from the bronchi, lungs were homogenized using the gentleMACS dissociator from Miltenyi Biotec (Auburn, California), diluted in phosphate buffered saline, centrifuged at 250 rpm for 10 minutes, treated with aqueous ammonium chloride to remove host cells, and used to prepare slides for enumeration [11].

### Microscopic enumeration

Slides were made by dropping 10μl of the homogenized lungs onto glass microscope slides within a circumscribed area and allowed to air dry. The slides were heat fixed and stained with Cresyl-Echt Violet (CEV) (Fisher Scientific) for enumeration of asci and rapid Wright-Giemsa (Fisher Scientific) for enumeration of all life cycle stages. *P. murina* asci and nuclei were enumerated by counting 30 microscopic fields at 1,250x power [13]. Data were expressed as log_10_ mean ± standard deviation per lung. GraphPad Prism v. 5 [12] was used to determine significance using the unpaired *t-*test.

### Survival Curve

At each time point or depending on the health of the mouse, 8 mice were sacrificed per group. The survival of each group was analyzed by GraphPad Prism v.5 using the Log-rank (Mantel-Cox) test and the Gehan-Breslow-Wilcoxon test. We typically stop these experiments at or around 8 weeks due to the poor condition of the mice and humanely sacrifice them at this endpoint.

## 3. Results and Discussion

Male and females in 3 mouse strains were evaluated for Pneumocystis burden at 3-time points that represent early, mid- and late infection (Figures 1 and 2). In Figure 1, the data were parsed to compare fungal burdens in male and female groups within each strain. Significant differences were observed in total lung burden (Fig. 1A, “Nuclei Counts”) at 6 weeks for all gender pairs with females carrying larger burdens. After 8 weeks of immunosuppression, the total organism burdens were significantly higher in the C3H/HeN and the C57BL/6 female mice. No significant differences were observed in the total lung burdens at the early (4week) time point. Interestingly, only a single significant gender difference was observed in asci numbers between male and female mice and that was at 6 weeks in the C3H/HeN strain (Figure 1B, “Asci Counts”). Since the Nuclei Counts represent all life cycle stages including the asci, these results may indicate an increased trophic burden in females vs males in the other 2 strains.

**Figure 1.**
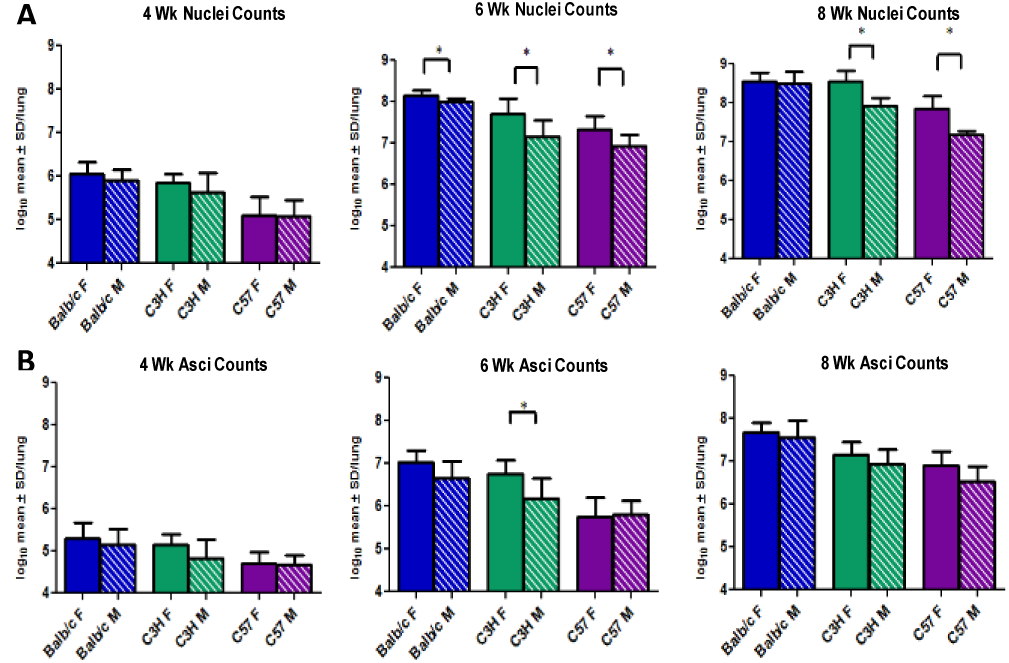
*Pneumocystis murina* burdens of males and females in 3 mouse strains. At 4, 6, and 8-week asci and nuclei (total fungal burden) of male (M) and female (F) Balb/c, C3H/HeN (C3H), and C57BL/6 (C57) were quantified by microscopic enumeration, log transformed, and expressed as the log_10_mean ± the standard deviation per lung (Y-axis). (*) Indicates statistical significance at p<0.05 using an unpaired *t-*test between male and female m of the same strain.

**Figure 2.**
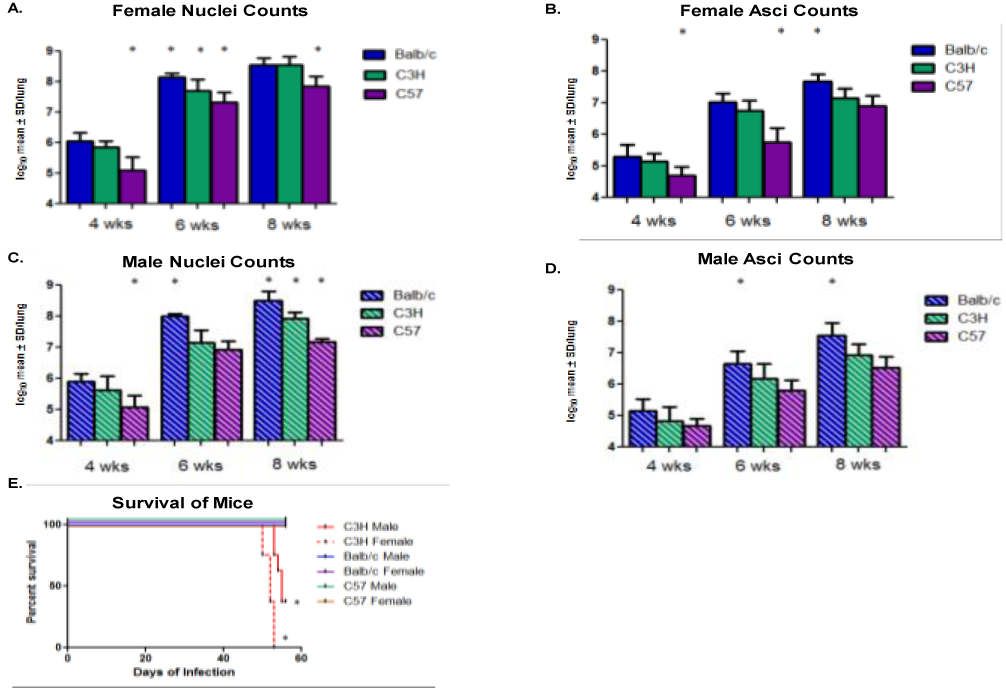
Strain and gender differences in burden and survival. The data presented in Figure 1 were parsed for gender and strain differences. (A) <u>Female nucle</u>i <u>count</u>. * indicates statistical significance of C57 mice nuclei burdens at 4 and 8 weeks compared to the other strains. (B) <u>Female asci coun</u>t. * indicates statistical significance of asci burdens of female C57 mice compared to the other strains at 4 and 6 weeks. (C) <u>Male nuclei count.</u> * indicates statistically significant differences in organism burdens. (D) <u>Male asci coun</u>t. * indicates statistically significant asci burdens of Balb/c mice when compared to other strains at 6 and 8 weeks. All statistical significance was set at p<0.05 using One-Way ANOVA and Newman-Keuls Multiple Comparison Test. Survival curves for all mice strains. (E) Survival curves were analyzed using GraphPad Prism v.5. (*) Indicates statistical significance at p<0.0088 for the male C3H/HeN and P<0.0001 for the female C3H/HeN.

Next, the P. murina burdens were compared within the same gender in the different mouse strains (Figure 2). In terms of total lung burden, the C57BL/6 female mice had significantly lower burdens at all 3-time points compared to females in the other 2 strains (Figure 2A). The Balb/c and C3H/HeN female mice had a significant difference only at 6 weeks (Figure 2A). Few differences were notable for numbers of asci in the female mice across strains (Figure 2B), apart from lower numbers in the C57BL/6 mice at 4 and 6 weeks, and at 8 weeks where the BAlb/c females had higher numbers of asci.

Analyzing the total lung burdens in male mice across strains, males of the C57BL/6 strain had burdens significantly lower at the 4 and 8-week time points, like their female counterparts (Figure 2C). Although burdens at 6 weeks in this group were lower than those of males in the other strains, significance was not reached (Figure 2C). The total fungal burdens in Balb/c male mice were significantly higher than males of the other 2 strains at the 6 and 8-week time point (Figure 2C). In the male mice, asci burdens were statistically different in the BAlb/c mice at 6- and 8 weeks, where again, there were higher numbers than in the other 2 strains (Figure 2D).

The males and females in the C3H/HeN strain experienced a higher mortality rate than the other strains (46 vs 100%) (Figure 2E), with all female mice succumbing to the infection or which had to be sacrificed due to morbidity prior to the end of the experiment. Note, in this animal model, all mice are sacrificed at the terminal endpoint of 60 days due to morbidity associated with the infection. Survival curves are based on mice who expired prior to the endpoint.

Adhering to NIH sex inclusion policy, we showed that gender did influence the progression of *P. murina* infection in this study, with higher fungal burdens observed in females of all 3 mouse strains at the mid-time (6 weeks) point and for C3H/HeN and C57/BL6 female mice after 8 weeks of immunosuppression. Reasons for this gender disparity could relate to factors in the lung environment, differential responses to the immunosuppressive regimen, or in the immune response to these fungal pathogens. It is interesting to note; few differences were observed in the numbers of asci regarding strain or gender. Balb/c male or female mice were the most permissive to this fungal infection as evidenced by the highest lung burdens. The outcomes from this study should guide future experimental design in pathogenesis of infectious diseases; systems biology “omics” approaches; and computational models of disease progression. The bases of these gender differences should be explored in future studies. Notably, these findings reiterate the importance of examining the effects of disease progression in both sexes, since such differences could impact clinical response to anti-Pneumocystis therapy or indicate distinct pathologies.

## Funding Acknowledgement

This work was supported by National Institutes of Health at the National Heart, Lung, and Blood Institute R01 HL1119190 and Administrative Supplement for Research on Sex/Gender Differences; the Veterans Affairs Biomedical Research Program I01BX000523.

None of the authors have a conflict of interest with any information in this report

This information has in part been presented at the International Workshops on Opportunistic Protists 13 (Seville, Spain 2014) and 14 (Cincinnati, OH 2017)

